# Endodermal suberin deposition restricts potassium leakage from roots

**DOI:** 10.1101/2022.12.20.521223

**Authors:** Morten Winther Vestenaa, Søren Husted, Francesco Minutello, Daniel Pergament Persson

## Abstract

- The endodermis is a checkpoint for ions and water escaping or entering the root. It has been hypothesized that suberin acts as a physical barrier preventing potassium (K) leakage from the stele during translocation, but attempts to support this idea has yielded contradictory results.
- We developed a Laser Ablation-Inductively Coupled Plasma-Mass Spectrometry (LA-ICP-MS) based element bioimaging method to study K leakage from roots with different suberin deposition, where we show that cesium (Cs) is an excellent tracer for K.
- Element bioimaging of roots and total shoot concentrations from various *Arabidopsis thaliana* mutants all showed a positive relationship between suberin deposition and K translocation efficiency. In addition, images from the fully suberized barley (*Hordeum vulgare*) seminal roots revealed a strongly reduced K leakage compared to less suberized root zones.
- Nodal roots form a scattered deposition of suberin towards the phloem in the mature root zone. This incomplete suberin deposition also restrict K leakage efficiently.
- Collectively, our findings provide experimental evidence that suberin act as a barrier for K leakage upon root-to-shoot translocation by restricting K movement over the endodermis from the stele to cortex.

## INTRODUCTION

Plants has developed roots with specialised tissues to acquire nutrients and water to support growth of the shoot (Hetherington & Dolan, 2018). Roots are often referred to as the hidden half of plants, and many questions related to their basic functions remain unresolved.

Between the vascular tissue and the surrounding cortex tissue is a single cell layer called the endodermis, where a physical barrier of suberin is deposited. Suberin monomers are synthesised in the endoplasmic reticulum (ER) and transported in vesicles to the extracellular space, where it is polymerized to suberin and deposited between the plasma membrane and the primary cell wall (Enstone *et al*., 2002; De Bellis *et al*., 2022). Suberin is highly hydrophobic and allow plants to prevent and alleviate various biotic and abiotic stresses. Accordingly, enhanced or delayed suberin deposition in roots are well-documented responses to e.g. salt stress, nutrient deficiency and toxicity, pathogen attack and drought. (Baxter *et al*., 2009; Krishnamurthy *et al*., 2009; Kreszies et al., 2018; Chen et al., 2019; Holbein et al., 2019).

Deposition of suberin varies in a gradient along the root axis with un-suberized cells at the root apex to complete suberization of all endodermal cells in the mature zones of the root. However, the complete suberized zone can be interrupted by single cells lacking suberin deposition, designated as passage cells. The gradient of increased suberin deposition upwards appears in three distinct maturation zones categorized as the unsuberized zone, a patchy zone, and a fully suberized zone (Grube Andersen et al., 2018). In barley, a fourth, so-called phloem-facing zone has been described, where suberin exclusively deposits in endodermal cells facing the phloem poles, thus, leaving all endodermal cells facing the xylem poles unsuberized (Chen et al. 2019).

The proposed role of suberin in regulating nutrient uptake remains enigmatic, but it has been proposed to restrict access of ions to plasma membrane nutrient transporters and control their expression in cells associated to the endodermis (Barberon & Geldner, 2014; Grube Andersen et al., 2018). Suberin deposits adjacent to the outer plasma membrane of endodermal cells where it obstructs access to transmembrane nutrient transporters that facilitate inward transport into endodermal cells. Suberin is, however, hypothesised to have dual functions to the root, as it both restricts backflow of nutrients out of the stele at the pericycle and endodermis interface and restricts ion influx by hindering transport at the boundary between endodermis and cortex (Barberon et al., 2016). Additionally, suberin likely have a secondary impact on nutrient uptake by restricting expression of nutrient transporters, as demonstrated for the phosphate transporter PHO1 (Grube Andersen et al., 2018). In Andersen et al. (2018), a funnel-like pattern of PHO1 expression was observed in the cortex and stele tissue adjacent to passage cells lacking suberin, indicating that suberin might restricts signaling leading to activation of nutrient transport.

Prior to the identification of suitable suberin mutants, the evidence collected on the role of suberin in nutrient uptake was extensive, but merely correlative (Schreiber et al., 2012). Increased suberin deposition from the tip to the base coincides with a general maturation of the root. Therefore, there is a significant risk that the impact of suberin on nutrient uptake, in the body of studies before suberin mutants became accessible, rather reflects changes in nutrient transporter activity, root morphology or other parameters associated with root maturation, than direct effects from suberization. Recent advances in the understanding of suberin and its relation to plant nutrition has come from studies comparing plant nutrient transporter- and suberin mutants. The combined studies of these *Arabidopsis thaliana* mutants have enabled researchers to isolate the role of suberin with respect to uptake and translocation of nutrients. Thus, our current understanding of how suberin influences ion translocation mainly comes from ionomic root and shoot profiles of mutants displaying no, decreased or ectopic suberin deposition (Baxter et al., 2009; Naseer et al., 2012; Li et al., 2017; Cohen et al., 2020). However, the collective evidence gathered from studying suberin mutants is still not conclusive regarding the role of suberin in nutrient uptake. For example, shoot concentrations of phosphorous (P), K and rubidium (Rb) decreased in the ELTP::CDEF1 and CASP1::CDEF1, i.e. mutants with unsuberized barriers, whereas the concentrations of the same nutrients increased in the shoot of the sub mutant, which also have a significantly decreased suberin deposition (Barberon, et al., 2016; Cohen et al., 2020). Additionally, mutants without suberin consistently show either no or only minor ionomic changes compared to wild types (Barberon et al., 2016). However, a possible explanation to this lack of clear shoot phenotypes, might be that plants compensate for a compromised suberin barrier by increased expression and/or activation of nutrient and water transporters (Barberon et al., 2016; Cohen et al., 2020). This is especially relevant for K mutants, considering that changes in K^+^ shoot concentrations is the most frequent nutritional phenotype of suberin mutants and that K^+^ play an essential role in stomata function and water relations as an osmotic regulator (Farooq et al., 2009). Accordingly, a critical milestone in the pursuit of understanding the role of suberin in K translocation is to experimentally overcome the fact that suberin is multifunctional, i.e. functions both as a barrier to nutrients as well as to water (Baxter et al., 2009; Barberon & Geldner, 2014).

As suberin seals the stele and thereby restricts K^+^ leakage, it facilitates efficient K^+^ translocation by restricting K^+^ loss rather than being a barrier for K^+^ uptake. This theory explains why an increased suberin deposition and an increased barrier function facilitates increased K^+^ translocation in roots. Moreover, the uniquely high mobility of K^+^ in plants also supports this idea. In contrast to e.g. Ca, Mn and B, K^+^ flows “freely” in the xylem, phloem and cytoplasm since it does not bind strongly to plant cell walls, ligands or proteins. Because K^+^ flows freely in plant tissue, it is suggested to be prone to leakage out of the stele during translocation (Barberon et al., 2016; Persson et al., 2016). Accordingly, it is likely that enhanced suberin deposition increase the K^+^ net uptake primarily by reducing loss and not by obstructing its inwards movement towards the vasculature of the root (Persson *et al*., 2016; Doblas *et al*., 2017).

We have developed a LA-ICP-MS method with cellular resolution, allowing us to image the lateral K distribution in root tissue sections of Arabidopsis mutants and barley roots differing in their suberization patterns. These analyses allowed for a detailed study on how suberin deposition influence K leakage in different root zones. We found that the degree of suberization was strongly correlated with K translocation efficiency, both in Arabidopsis mutants and seminal roots of barley. Accordingly, we saw that fully suberized root zones strongly reduced the K leakage over the endodermis, compared to zones with less suberization. However, nodal roots of barley appeared to constitute a special case since they also effectively restricted K leakage despite their incomplete “phloem facing” suberin deposition. We propose that while suberin physically blocks the loss of K at the suberized endodermal cells facing the phloem, the activity of K transporters in passage cells and associated cells facing the xylem poles are able to rescue K that would otherwise be lost.

## MATERIALS AND METHODS

### Plant Material and Experimental Setup

Seeds of barley (*Hordeum vulgare L*. cv. Irina) were germinated for 7 days in vermiculite followed by cultivation in a greenhouse with minimum day/night temperatures of 18/15 °C, humidity 60% and a day/night cycle of 16 h/ 8 h, using supplementary light of 300 μmol photons m^-2^ s^-1^ by high-pressure sodium bulbs. After germination, seedlings were transferred to aerated hydroponic units containing 4L nutrient solution with the following composition: 200 μM KH_2_PO_4_, 200 μM K_2_SO_4_, 300 μM MgSO_4_·7H_2_O, 100 μM of NaCl, 300 μM Mg(NO_3_)2·6H_2_O, 900 μM Ca(NO_3_)24·H_2_O, 600 μM KNO_3_, 50 μM Fe(III)-EDTA Na, 1 μM MnCl·4H_2_O, 0.8 μM Na_2_M·O_4_·2H_2_O, 0.7 μM ZnCl_2_, 0.8 μM of CuSO_2_·5H_2_O, 0.8 μM NiCl_2_, 2 μM H_3_BO_4_ and 40 μM Na_2_Si_2_O_2_. All nutrient solutions were prepared in Milli-Q water (Milli-Q Element; Millipore). The nutrient solutions were changed weekly and the pH in each hydroponic unit was maintained at 6.0 ± 0.3 using ultrapure HCl.

### K uptake in Nodal and Seminal Roots

A split-root experiment was performed to investigate the translocation efficiency of seminal and nodal roots of barley, respectively. After two weeks in hydroponics, the plants were moved to 250mL blue cap bottles wrapped in aluminum foil, with nodal and seminal roots gently divided into separate bottles; one root type per bottle. The bottles of the treated plants (n=4) contained a modified nutrient solution containing 0.1 mM 95.80% enriched ^41^K isotope (Trace Science International, USA) as sole K source, where either nodal or seminal roots were given the ^41^K isotope separately. Following 6 hours of K isotope incubation, the youngest fully emerged leaf (YFEL) of each plant was harvested for stable isotope ratio analysis by ICP-MS, including the YFELs of untreated plants. The amount of translocated isotope in the treated plants was calculated as the ^41^K enrichment relative to the untreated controls (Δ^41^K_treated root type_-^41^K_control plant_), per root per unit time of exposure to ^41^K (Figure 2).

### Element analysis of leaves

All leaf samples were digested using an UltraWAVE Pressurized Microwave Digestion System (Anton Paar GmbH, Austria) with 2.5 mL of HNO_3_ (70%, v/v) and 1 mL of H_2_O_2_ (15%, v/v), and subsequently diluted to 50 mL, resulting in a final concentration of 3.5% HNO_3_ (Hansen et al., 2013). For stable isotope ratio analysis, element analysis was performed on an ICP-MS (8900 Triple Quadruple ICP-MS, Agilent Technologies, USA) in H_2_ mode (6.5 ml/min) with optimised spectrum recording mode for isotope ratio (1 point peak pattern, 1 replica and 10.000 sweeps per replica). The ion intensity was recorded at mass 41 as counts per second (cps) and statistically analysed by a student’s t-test in Sigma Plot. Element concentration analysis of Cs in rosettes and barley shoots was performed on an ICP-MS (8900 Triple Quadruple ICP-MS, Agilent Technologies, USA) operated in standard mode. Tissue concentrations were calculated in MassHunter (Agilent Technologies, USA), and statistically analysed by a one-way ANOVA in Sigma Plot14.

### Cesium Leakage during Root to Shoot translocation

Three *Arabidopsis thaliana* mutants (esb1, CASP1:CDF1 and wild type Col-0) were cultivated in hydroponics according to (Lilay et al., 2019). Briefly, plants were grown in 1.5 mL microtubes placed in lids of 10L plastic containers containing a modified half-strength MES-buffered Hoagland solution. Plants were cultivated in a climate chamber using a 16/8 hours light/dark cycle with 22/20°C light/dark temperature, 70% humidity and 125 μmol photons m^-2^ s^-1^ provided by light emitting diodes (Senmatic FL300 Sunlight). Each individual plant was removed from the hydroponics, then root tips were exposed to 25 μL of a 10 mM CsCl solution for 30 minutes. Subsequently, rosettes and 1 cm sections from the main axis of the primary roots were sampled at 75% distance from the root tip. The rosettes were prepared for element concentration analysis by ICP-MS, whereas the root segments were prepared for LA-ICP-MS based multi-element imaging, as described below.

### Histological Analysis of Root Endodermal Suberin

Staining of suberin in intact, whole roots was performed in order to determine the longitudal suberin patterns (Chen et al. (2019)). Whole roots were fixed with 4% paraformaldehyde, using vacuum infiltration for one hour and then cleared in ClearSee solution (100 g/L xylitol 150 g/L sodium deoxycholate and 150 g/L urea (Kurihara et al., 2015)) for one week prior to staining by Fluorol yellow (FY). Root sections were stained using a fresh solution of 0.01 % w/v FY in lactic acid (1 hour at 70°C) and then post-stained for 1 hour with 0.5% w/v aniline blue. Stained roots were cut in 5 cm long pieces and placed in a few drops of 50% glycerol on glass slides. Seminal roots were observed with a fluorescence microscope at 460/40 nm emission and 510/30 nm excitation (DMR HC, Leica, Germany), but due to their relatively large diameter, nodal roots were observed in a confocal laser-scanning microscope (Stellaris 8, Leica Germany) with 475 nm excitation and 500-520 nm emission. Images presented in figure 1 are 3D renderings of z-stacks obtained on this confocal laser-scanning microscope. Three distinct zones were identified and recorded along the root axis, according to differences in their suberization patterns. In all seminal roots, these zones were identified as “unsuberized zone”, “patchy zone”, “phloem-facing zone” and “mature zone”. These zones were identified in order to quantify the extent of the suberin layer development in each root. In the nodal roots, only the “unsuberized”, “patchy zone” and the “phloem-facing” zone was found. The extent of each suberization zone in the approximately 30 cm long roots, was normalized to total root length and reported as the percent fraction of the total root length.

**Figure 1.**
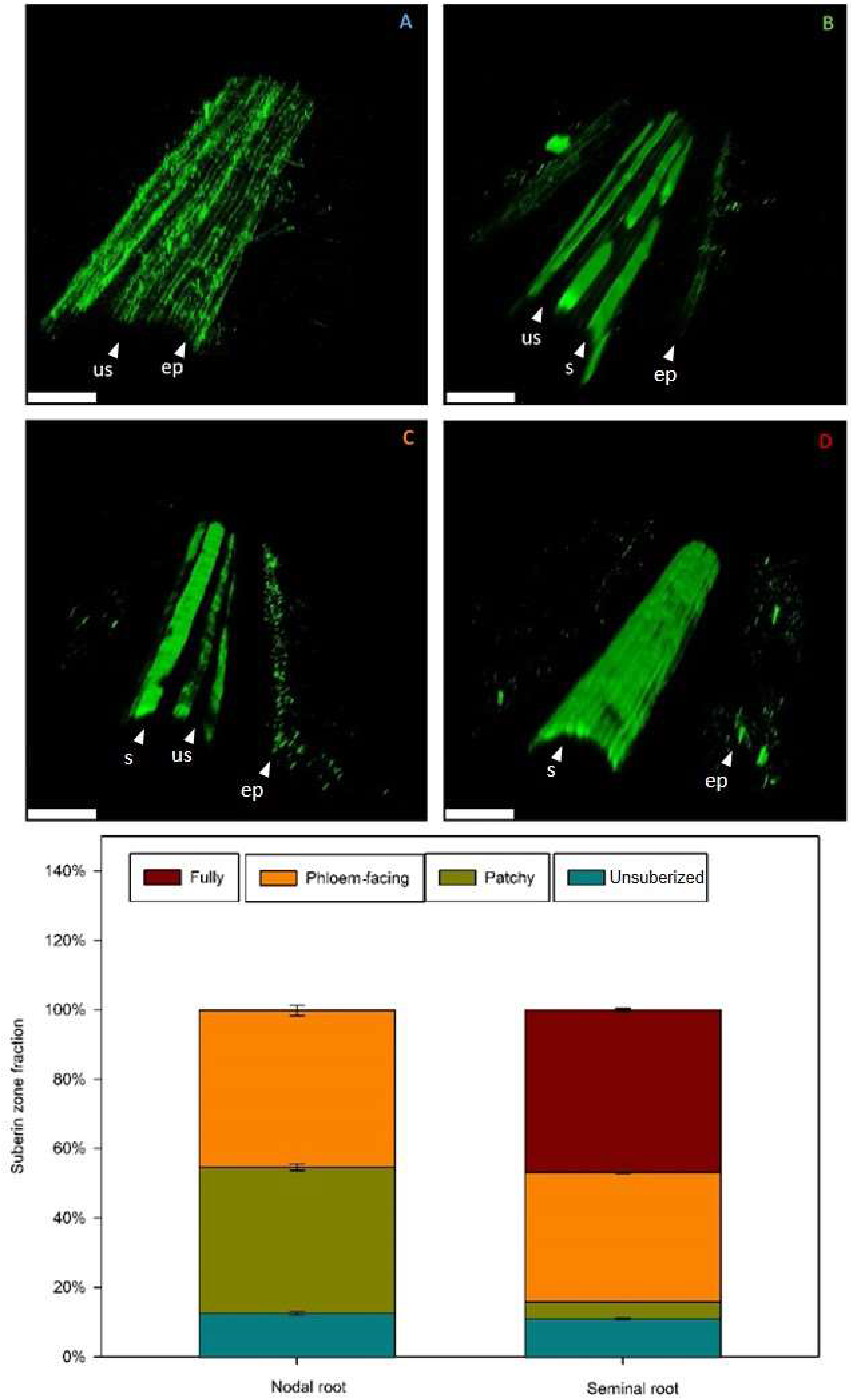
The endodermal suberin deposition along the nodal and seminal roots of barley (*Hordeum vulgare*). Suberin deposition was observed by fluorescence microscopy in longitudinal sections by clearing with ClearSee and then staining by fluorol yellow (FY), using roots of identical physiological age (approx. 30 cm total root length). The green signal in endodermis indicate FY stained suberin, whereas green signal in epidermis indicate autofluorescence with considerably lower intensity, possibly originating from lipids associated to root hairs. Arrows indicate epidermis (ep), suberized cells (s) and unsuberized cells (us). A, B, C and D represent the four distinct suberin zones observed in seminal roots. At the root apex an unsuberized zone was found (A); followed by a zone with patchy suberin (B) and a phloem-facing zone (C) which is characterized by unbroken longitudinally rows of suberized cells. Finally, the phloem-facing zone develops into a fully suberized zone (D). Scalebars are 200 μm. (E), Suberin zones of seminal and nodal roots, reported as relative fractions of the total root length in percentage (cm suberized root length per total root length), measured from the root apex. The fully suberized zone was always lacking in nodal roots, where instead the phloem-facing zone continued all the way to the base of the root. Error bars represent one SE (n=4).

### Multi-element Imaging by Laser Ablation Inductively Coupled Plasma Mass Spectrometry

Multi-element imaging was performed on a Laser Ablation Inductively Coupled Plasma Mass Spectrometry system (LA-ICP-MS) using a 193 nm ArF excimer laser (Iridia, Teledyne Photon Machines, Thousand Oaks, CA 91360, USA) equipped with a Cobalt cell (Teledyne Photon Machines) and connected to an Inductively Coupled Mass Spectrometer (model 8900, Agilent Technologies, USA). Analysis was done with the following settings: Scan speed: 160 μm s^-1^, Repetition rate: 200 Hz, Spot size: 4 μm circles for the images in figure 5, 6 and 7, and fluence: 0.5 J cm^-2^. Helium (He) was used as transfer gas from the laser ablation unit to the ICP-MS with a flowrate of 0.15 mL min^-1^ and the ICP-MS was operated in standard mode with a sample cone depth of 30 mm and a nebulizer gas flow (Ar) of 0.75 mL min^-1^. The Mass Spectrometer was set to scan across the following masses ^24^Mg, ^39^K, and ^133^Cs, in a scan cycle of 25 ms using integration times of 6 ms for ^24^Mg, 1 ms for ^39^K and 10 ms for ^133^Cs. Any drift in sensitivity was monitored before, between and after all analyses, using a NIST612 glass standard (NIST; National Institute for Standards and Technology, SC, USA). All images were processed with the HDIP (High Definition Image Processing) software (HDIP, Teledyne CETAC Technologies) including gas blanks and signal drift corrections.

### Sample Preparation for Multi-element Imaging

Samples for LA-ICP-MS were prepared according to Persson *et al*., (2016), describing a method that preserves the native ion environment as well as the cellular structures of root cross sections. Briefly, an approximately 1 cm root segment was sampled, and all lateral roots were removed from this segment with a scalpel. Hereafter, the root sample was briefly dried on a napkin, then encapsulated in melted paraffin and placed in an aluminium foil mold filled with OCT medium (Optimal Cutting Temperature, Tissue-Tek from Sakura Finetek, Japan). The mold with the encapsulated root sample was then snap-frozen in liquid N_2_ and stored in a −20°C freezer until sectioning. For sectioning, the OCT-blocks with root samples were transferred to a pre-cooled (25°C) cryotome (Leica model CM050S) and were left to equilibrate for 1 hour. The first ~2 mm of the root segment were discarded, then 16 μm thick cross sections were cut and transferred to cold microscope slides (SuperFrost Plus, VWR International using cryofilm (2C, Section-lab Co. Ltd, Japan) according to the Kawamoto cryofilm method (Kawamoto & Kawamoto, 2014). Microscope slides with root cross sections were transferred to a −20°C freezer to freeze-dry overnight. Sections were evaluated using bright field microscopy and the sections of highest quality was selected for analysis (DM2000 microscope with FLEXACAM C1 mounted using LAS X Life imaging software, Leica, Germany).

### Sampling for Multi-Element imaging

It is not possible to image any of the K isotopes using LA-ICP-MS due to the high natural abundance of K in the samples. Thus, experiments were instead performed using the K proxy ^133^Cs^+^, which display the same physicochemical properties as K^+^ and is non-toxic when used in sufficiently low concentrations. In order to investigate and document outward radial ion movement over the endodermal suberin barriers, root segments of living plants were isolated, treated and then sampled based on *a priori* knowledge to the distribution pattern of suberin along the longitudinal axis of seminal and nodal roots. Treatment and sampling was performed at 25% and 75% distance from the root tips of both root systems.

The plant to be treated was mounted with the shoot upwards and with the root placed horizontally, on moist paper, in a plastic box. The area to be treated was localized using a ruler and then a 1 cm segment on the living root was isolated from the rest of the root with grease barriers, approx. 1 cm apart from each other, which was manually applied to the root (Grease for Pivots Cat. No. 5810350.050-01, Eppendorf, Germany). On test plants, we used a 0.5% solution of Aniline Blue to document that the grease barrier efficiently stopped the solution from spreading along the surface of the root axis (data not shown). The two grease barriers thus defined the 1 cm root segment to be treated with the Cs proxy solution. We saturated a cotton ball with the proxy solution and placed the cotton in close contact with the root, along the entire 1 cm segment (with a grease barrier both upwards and downwards). The Cs solution was supplied to each isolated root segment for 10 min and then an unexposed root segment (1 cm) upwards from the upper grease barrier (towards the shoot) was harvested. The idea was to apply Cs and have it move radially towards and into the stele in the treated section, then upwards towards the shoot, while still in the stele. This set-up enabled us to study possible ion leak outwards from the stele during upward translocation. Samples were also harvested 2 cm below the grease barrier, which served as controls to document that Cs had not escaped from the lower the grease barrier. As no Cs could be detected in these root segments, we are confident that the Cs signals recorded in the samples had exclusively entered the plant radially, in the cotton-treated area.

## RESULTS

### Suberin deposition in the endodermis of barley nodal and seminal roots

Four root zones with distinct suberin patterns were identified along the main axis of seminal roots, but only three zones were found in the nodal roots (Fig. 1, E). The endodermis did not contain suberized cells within the initial 15% of nodal and 10% of seminal roots, measured from the root apex (Fig. 1, D and E). In accordance with Chen et al. (2019) this zone was designated as the unsuberized zone. Following this zone, a zone characterised by single and scattered suberized endodermal cells was observed, which was designated as the patchy zone. In nodal roots, this zone appeared from 15 to 55% distance from the root apex, but only from 10 to 15% in the seminal roots (Fig. 1, C and E). Further towards the root base, we observed a pattern of uninterrupted lines of suberized cells. Chen *et al*. (2019) coined this zone the phloem-facing zone, since the suberized endodermal cells in this zone were exclusively facing the phloem poles of the stele, thus leaving lines of unsuberized cells facing the xylem poles (Fig. 1, B and E). These xylem-facing, unsuberized cells have previously been designated as passage cells due to lack of suberin (Geldner, 2013; Holbein *et al*., 2021). We found that the phloem-facing zone covered the entire nodal root axis from the end of the patchy zone to the root base, thus corresponding to the uppermost 55-100% of this root type (Fig. 1, B and E). In contrast, the phloem-facing zone of seminal roots only appeared at a 15-55% distance from the root apex and was followed by a zone where suberin covered all but a few, randomly distributed passage cells. This zone, which was exclusively found in seminal roots, was designated the fully suberized zone and covered the endodermis at a 55-100% distance from the root apex (Fig. 1, A and E).

### Net assimilation of Potassium in leaves taken up by seminal and nodal roots

The K uptake capacity of nodal and seminal roots, respectively, was quantified by analysing the enrichment of YFELs with the stable ^41^K isotope (Fig. 2). The ^41^K isotope solution was applied to either the nodal or the seminal root system of each plant in a split-root setup, and 6 hours after isotope exposure the YFELs were harvested and a significant enrichment was observed in the shoot relative to the untreated controls. On a root and time basis, the ^41^K ion counts measured in the leaves increased by 8% when supplied to seminal roots, and by 10% when supplied to nodal roots. Statistical analysis showed an insignificant difference in enrichment of ^41^K in YFEL between nodal and seminal root systems (p=0.2). ^41^K enrichment were not corrected for root DW as there was no difference in root DW of nodal (p=0.52) nor seminal (p=0.63) roots between treatments.

**Figure 2.**
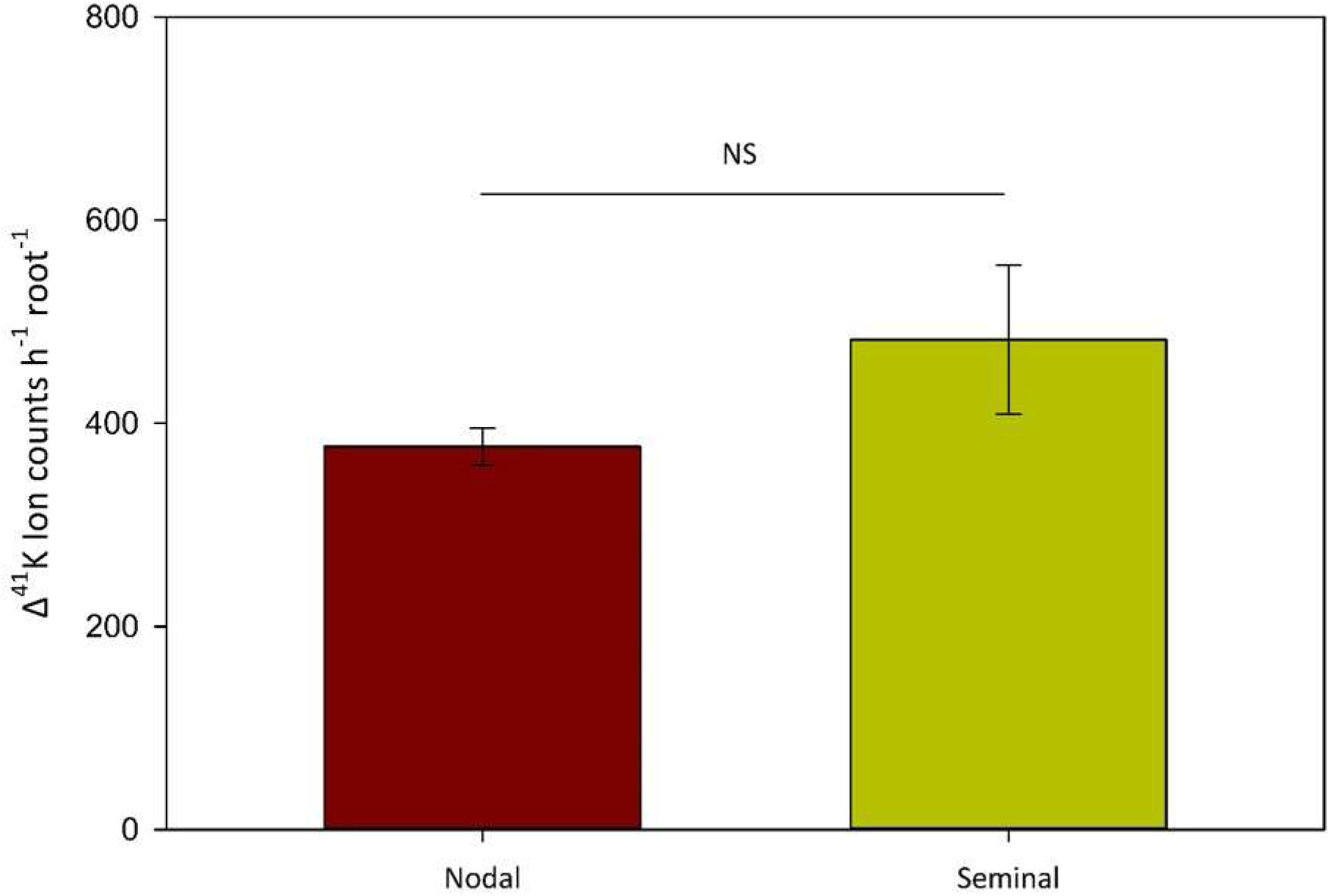
Change in ^41^K isotope ion counts in the youngest fully emerged leaves of barley (*Hordeum vulgare*), following 6 hours of ^41^K uptake and translocation by nodal or seminal root systems, in a split root setup. NS indicate an insignificant difference in uptake and translocation of the ^41^K isotope. Error bars represent one SE (n=4)

### Validation of Cs as a suitable tracer for bioimaging of K distribution in tissue

Using LA-ICP-MS, it was not possible to track changes in the tissue distribution of K by studying changes in one or more of the natural stable K isotopes, due to a combination of the high K background in root tissue and the relatively low analyte transfer efficiency from the Laser Ablation system. Instead, cesium (Cs) was implemented as a tracer for K, due to the physicochemical similarities between these two elements and due to the low background of Cs in plant tissue grown in hydroponics. The potential differences in Cs and K uptake and translocation patterns (Middleton et al., 1960) requires careful validation, hence it is of particular importance to document that the two ions co-localize in the tissue. Additionally, the potential risk of Cs toxicity requires that the applied Cs concentrations and exposure times are carefully optimized prior to root exposure. Indeed, no toxicity responses were observed in the concentrations and exposure times used in the following experiments. Root segments of nodal roots, sampled at a 75% distance from the root apex were incubated with 10 mM Cs for 10 minutes, then harvested and prepared for LA-ICP-MS analysis. In the resulting images, we observed a close correlation (R^2^ = 0.76) between the ion intensities of Cs and K across the root (Fig. 3A), whereas a significantly weaker correlation was observed between Cs and Mg (R^2^ = 0.44), showing that the biodistribution of Cs was largely decoupled from that of other essential macro nutrients such as Mg. In addition, a weaker Mg signal was found in the stele compared to that of the cortex, as opposed to the uniform distribution of both Cs and K between stele and cortex Fig.

**Figure 3.**
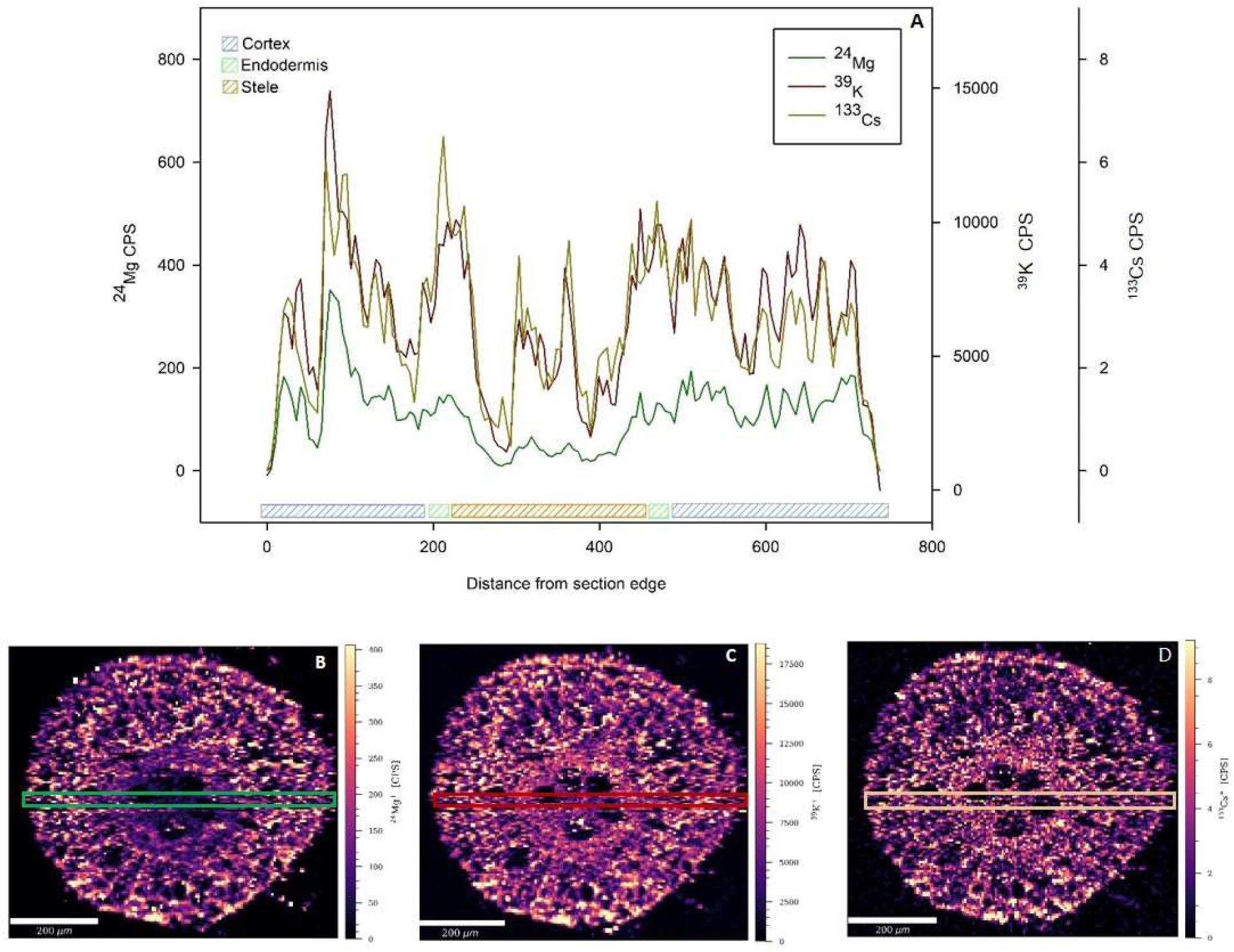
The distribution of ^24^Mg, ^39^K and ^133^Cs in a cross section from the mature part of a nodal barley (*Hordeum vulgare*) root, exposed to 10 mM CsCl for 10 minutes. A, the ion intensities in a line scan of ^24^Mg, ^39^K and ^133^Cs across the root section. Line scans were extracted at the broadest diameter of the root cross section and values are shown as means across 40 μm (spot size 4μm). The colored boxes represent localization of cortex, endodermis and stele in B, C and D; The tissue distribution of ^24^Mg (left), ^39^K (middle) and ^133^Cs (right), respectively, in the same root cross section, sampled at 75% total root length from the root apex. The sections were sampled within the incubated area and images represent intensities of ^24^Mg, ^39^K and ^133^Cs reported as counts per second (CPS). Line scans were analyzed in three independent biological replicates with similar results.

### Suberin deposition predicts root to shoot translocation efficiency of K

To further investigate if suberin restricts K leakage during upwards translocation in the root, we studied the root-to-shoot and tip-to-mature root translocation efficiency in three *Arabidopsis thaliana* mutants with contrasting suberin deposition, again utilizing Cs as a proxy for K. Following a 30 min addition of Cs exclusively to the root tip, leaf rosettes and root sections from the fully suberized zone were sampled from wild-type (Col-0) and suberin mutants (esb1, increased suberin; CASP1::CDEF1, decreased suberin). This enabled us to compare the effect of suberin on stele leakiness and the resulting translocation efficiency. Accordingly, the translocation of Cs resulted in mutant-specific Cs accumulation in the rosettes in the order esb1<Col-0<CASP1::CDEF1 (Fig. 4A) which clearly matches their relative suberin content (esb1>Col-0>CASP1::CDEF1) (Baxter et al., 2009; Naseer et al., 2012; Calvo-Polanco et al., 2021). These results confirm that suberin indeed restricts the loss of Cs, resulting in enhanced translocation towards the shoot. Cesium was accordingly only detected in the stele and cortex of esb1 roots at 75% distance from the root apex, with ion counts peaking at 1500 CPS (Fig. 4D). Thus, both in the Col-0 and in the low suberin mutant, a majority of the applied Cs was lost due to leakage during translocation from the root apex to the mature zone.

**Figure 4.**
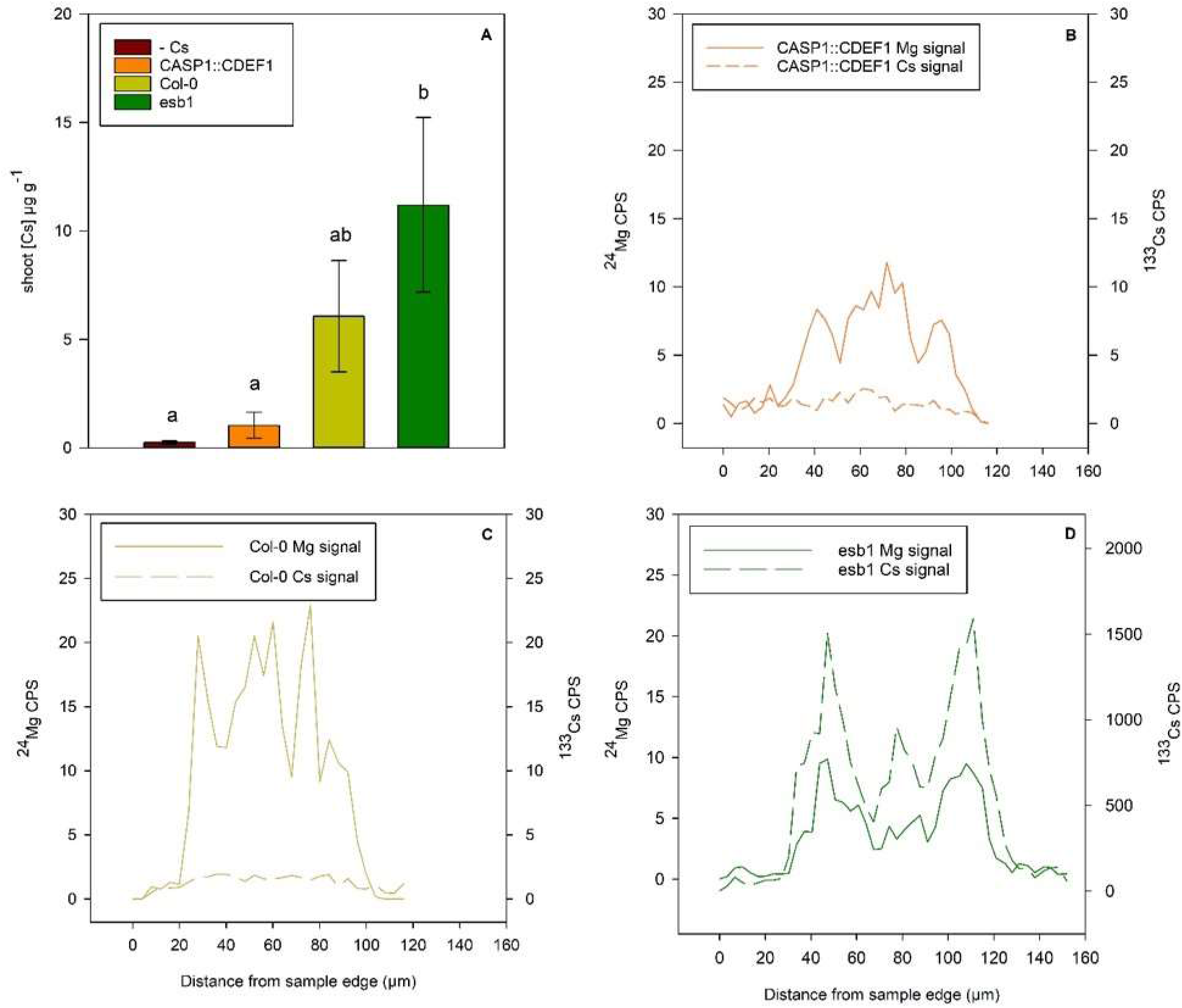
Upward translocation of cesium (Cs) in *Arabidopsis thaliana* mutants with contrasting endodermal suberin deposition. Root sections and rosettes were sampled following 30 min. incubation with 10 mM ^133^Cs, exclusively given to the root tip. A; Total shoot concentrations of Cs in Col-0, CASP1::CDEF1 and esb1 plants, respectively. Results in A are mean values ± SE (n=4). B, C and D, line scans of root cross sections sampled at 75% distance from root apex of Col-0 (C), CASP1::CDEF1 (B) and esb1 (D). Line scans of ^24^Mg and ^133^Cs from element images were extracted at the broadest diameter of root cross sections and the reported values are averages of the relative intensities across 10 μm (spot size 4μm). Line scans were analyzed in three independent biological replicates with similar results.

### Potassium leakage decreases in barley root sections with increasing suberin deposition

The endodermis varies in suberin deposition according to its maturation state (Fig. 1). As a consequence, ion leakage outwards across the endodermis could be analysed, by adding Cs to a series of isolated root segments differing in maturation state and then sampling the upwards, untreated root segments. Using this setup, it was observed that Cs is largely confined within the stele where the endodermis is fully suberized (Fig. 5A). Hence, a fully suberized endodermis restricts the radial leakage of K during upwards translocation, as indicated by the steep Cs gradient found at the interface between the endodermis and cortex tissues (Fig. 5, B). A contrasting Cs distribution was found in root sections with less suberin, sampled 25% from the root apex of seminal roots (Fig. 5C). Here, suberin only deposited around endodermal cells facing the phloem poles of the stele (Fig. 1), which resulted in a substantial leakage, as indicated by the homogenous radial distribution of Cs across stele and cortex (Fig. 5, C and D). In the nodal roots, a similar distribution of Cs was found in sections sampled at 25% distance from the root apex (Fig. 6, C and D), where only few cells were suberized, in a patchy pattern (Fig. 1, B). However, the phloem-facing zone of nodal roots appeared to restrict leakage from the stele, in a similar fashion as observed in the fully suberized zone of seminal roots. Accordingly, in nodal roots the steep Cs signal gradient between stele and cortex coincided with an endodermis suberized only in the phloem-facing pattern (Fig. 6, A and B). This observation show that an endodermis with suberized cells facing the phloem poles can efficiently restrict loss of K from the stele during upwards translocation, but only in the mature root tissues. In seminal roots, the phloem-facing zone is found closer to the apex, in less mature tissue, with a lower capacity to restrict radial K leakage.

**Figure 5.**
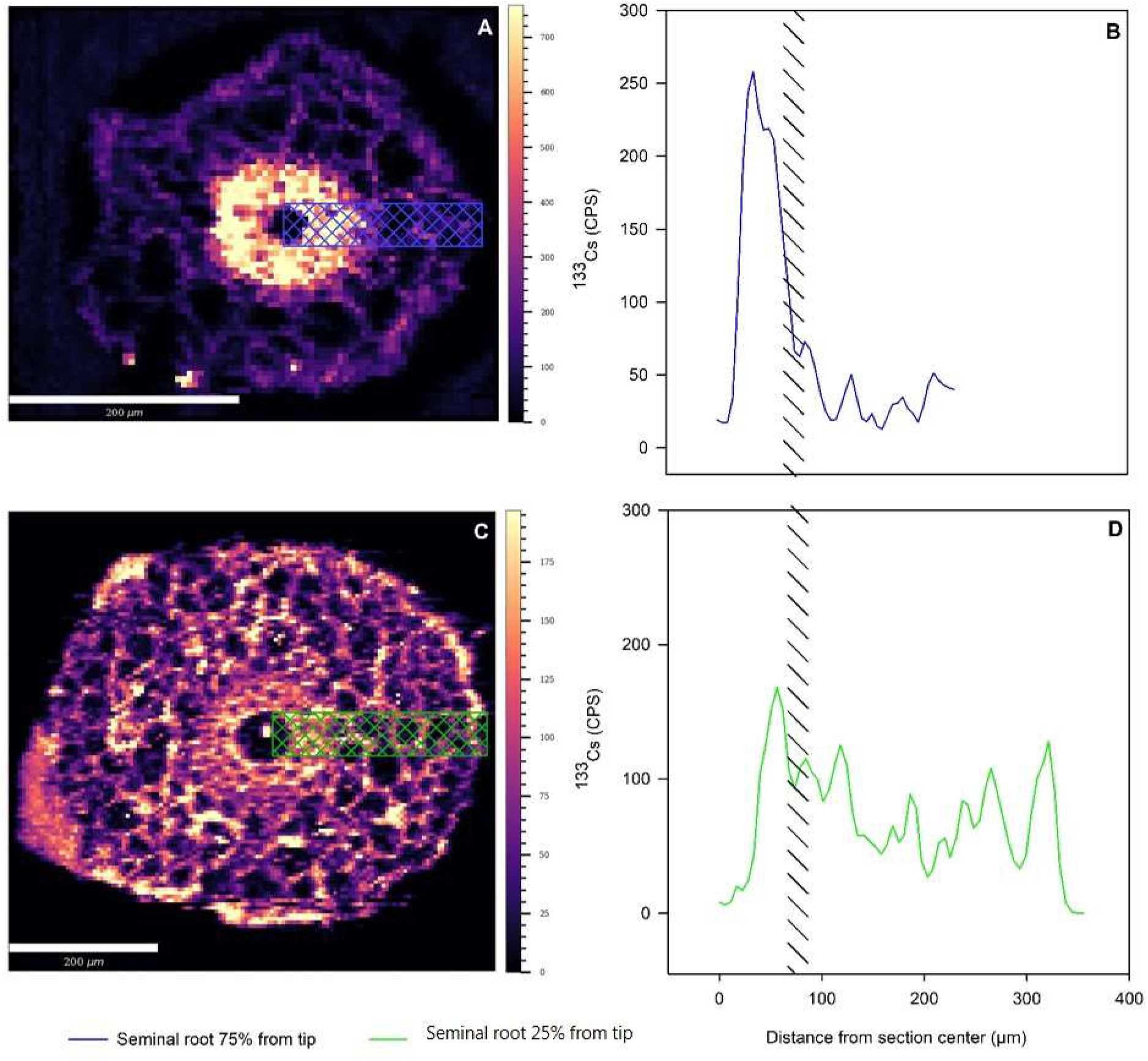
Cesium (Cs) distribution in the fully and phloem-facing suberized zones of barley (*Hordeum vulgare*) seminal roots. A and C, images of Cs distribution in root cross sections of root sampled in the fully suberized zone (A) and the phloem-facing zone (C) at distances equivalent to 75% and 25% of full root length from the root apex, respectively. Root sections were sampled following a 10 min. incubation with a 10 mM ^133^Cs solution. Samples were taken one cm upwards from the treated area; towards the root base, and images represent the ion intensities of ^133^Cs reported as counts per second (CPS). B and D, Line scans of Cs intensities across radial root cross sections A and C. We extracted line scans at the broadest diameter of root cross sections and reported values are average intensities across 40 μm. Colored crosshatched boxes in A and C represent line scans reported as relative ion intensities of ^133^Cs (CPS). Crosslines in B and D represent where the extracted line scan coincide with the endodermis. Line scans were analyzed in three independent biological replicates with similar results.

**Figure 6.**
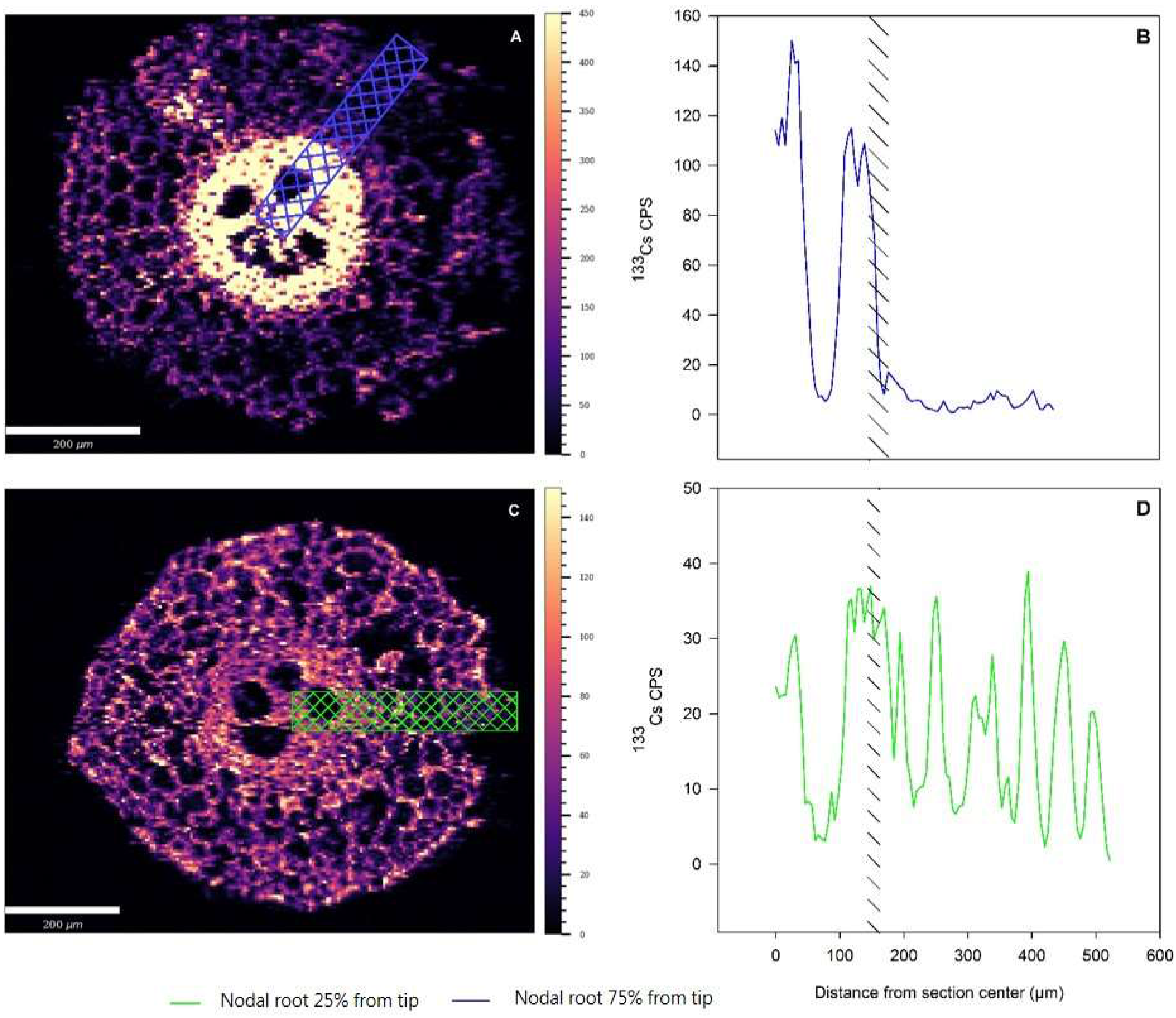
Cesium (Cs) distribution in the phloem facing and patchy-suberized zones of barley (*Hordeum vulgare*) nodal roots. A and C, Images of Cs distribution in root cross sections of root sampled in the phloem-facing zone (A) and un-suberized zone (C) at distances equivalent to 75% and 25% of full root length from the root apex, respectively. Root sections were sampled following a 10 min. incubation with a 10 mM ^133^Cs solution. Samples were taken one cm upwards of the treated area; towards the root base, and images represent ^133^Cs intensities reported as counts per second (CPS). B and D, line scans of Cs intensities across root cross sections in A and C. Line scans were extracted at the broadest diameter of root cross sections and values are reported as average ion intensities across 40 μm. Crosslines in B and D represent where the extracted line scan coincide with the endodermis. Line scans were analyzed in three independent biological replicates with similar results.

## DISCUSSION

Contrasting observations has previously been reported on the shoot ionomes of suberin mutants of *Arabidopsis thaliana* (Barberon *et al*., 2016; Cohen *et al*., 2020; Shukla *et al*., 2021; Calvo-Polanco *et al*., 2021). Accordingly, the effects of suberin on uptake and translocation of mineral ions remains enigmatic. In order to resolve the reason for these contrasting data, there is a need to investigate and clarify the mechanistic role of suberin in controlling uptake of specific nutrients. Although it is commonly agreed upon that suberin plays a role in nutrient uptake, data also suggest that suberin exert a range of additional, and maybe even more important functions in the roots. These additional functions include regulation of water flow, exemplified by the enhanced suberin deposition as a response to drought and salt stress, and the barrier functions towards pathogens (Baxter *et al*., 2009; Kreszies *et al*., 2018; Holbein *et al*., 2019; Calvo-Polanco *et al*., 2021). Consequently, there are no data available that unambiguously show, whether the suberin plasticity is a direct response related to the mineral nutrition of plants, or if its effect on mineral nutrition is a secondary response, originally related to e.g. water balance or other responses.

In the current study, we aim at presenting mechanistic evidence for the proposed role of suberin in K uptake and translocation in roots. It has previously been suggested that suberin support K net uptake by restricting K loss from the stele, by acting as a physical barrier for radial K movement across the endodermis (Barberon, et al. 2016). However, an experimental approach, studying the K dynamics across the suberin layer in root tissue did not exist prior to this work. Using LA-ICP-MS, we developed an experimental setup allowing us to image and mimic the distribution of K, using Cs as a proxy. This set-up allowed for imaging of the stele and cortex with cellular resolution, during root-to-shoot translocation. By imaging the distribution of Cs in various tissue regions and by analysing changes in radial Cs gradients across the endodermis, the role of suberin could be unravelled in root sections with contrasting suberization. Accordingly, we employed this technique to study two root types (seminal and nodal) of barley with contrasting suberin deposition (Fig. 1) and in three mutants of *Arabidopsis thaliana* with different suberin content; Col-0 (wild-type), CASP1::CDEF1 (reduced suberin) and esb1 (increased and ectopic suberin). The main findings are discussed below.

### Cs is a suitable K tracer for bioimaging by LA-ICP-MS

The ability of suberin to restrict radial leakage of K across the endodermis was initially studied by monitoring differences in K distribution between specific root tissues, using LA-ICP-MS based bioimaging. However, it is analytically very challenging to assess differences in K distribution, by merely studying changes in the K concentration at different time points, due to the high background concentration of K. Thus, analyzing the biodistribution of K requires the use of either K isotopes or suitable elemental proxies for K.

Radioactive isotopes are perfect tracers in terms of sensitivity, but such analyses typically produce a low image resolution, which hinders tissue-specific interpretation (Kopittke et al., 2020). Additionally, use of the radioactive isotope ^40^K in LA-ICP-MS is hampered by ^40^K being a beta-emitting radioisotope with a half-life of 1.3 × 10^9^ years. High radiance is not suitable for LA-ICP-MS imaging due to contamination of expensive lab equipment and safety issues. Stable isotopes are ideal tracers due their absolute similarity with the nutrient of interest. However, use of the stable potassium isotopes (^39^K; ^41^K) is not suitable for ICP-MS based imaging, due to a high natural background and the existence of overlapping polyatomic interferences, such as ^41^ArH, on ^41^K. Thus, relatively low amounts of ablated tissue using LA-ICP-MS, combined with interference of polyatomics and high analyte backgrounds makes detection of stable K isotopes impossible, even when using state-of-the-art equipment.

Due to the challenges using stable K isotopes, ^133^Cs and ^89^Rb are commonly employed tracers for K, since they belong to the alkali metals and predominantly exists as monovalent cations in solution, just like K. Rubidium (^89^Rb) is frequently used, but it is commonly observed as background contamination in plant tissue (Liu et al., 2020; Shukla et al., 2021). Rubidium is a contaminant in salts used to prepare hydroponic solution and accordingly high ^89^Rb background levels will always be found in plant tissue, preventing its use as a proxy for K. Cesium, however, is not a significant contaminant in the salts used for hydroponics, but may have toxic side-effects on plant growth (Hampton et al., 2004), and it has not previously been documented that Cs and K co-localise in plant tissue, at the cellular level. Here, Mg was included in the analysis as a control element, to show that the distribution of K and Cs is similar, on the tissue level. We show that Cs and K co-localises in root tissue, whereas Mg is uncorrelated to both K and Cs (Fig. 4). Furthermore, we have documented that there are no negative physiological responses to Cs at the concentrations and time of exposure used in this study (Supplementary Fig. 1).

### Suberin act as a barrier to prevent K leakage during root to shoot translocation

The efficiency of Cs translocation from root to shoot was found to be dependent on endodermal suberization (Fig. 4). Accordingly, mutants with ectopic suberin (esb1) had higher Cs concentrations in shoots than plants with unaltered suberin (WT), which is in accordance with Baxter et al. (2019), who previously reported increased shoot K concentration in the esb1 mutant. Furthermore, plants with unaltered suberin deposition in roots (WT) had higher Cs concentrations in their shoots than mutants lacking endodermal suberin; e.g. the CASP1::CDEF1 mutant (Fig. 4, A). The CASP1::CDEF1 mutant completely lacks suberin in the endodermis due to a constitutive expression of the Cutin Degrading Factor 1 enzyme (Naseer *et al*., 2012). The CASP1::CDEF1 results are unclear, as it has previously been reported to both lack a clear K phenotype (Calvo-Polanco *et al*., 2021) and to have a significantly decreased shoot K concentration (Barberon *et al*., 2016). However, Calvo-Polanco et al. (2021) only recorded a slight decrease in suberin content in the roots of the CASP1::CDEF1 mutant, which is in stark contrast to the observations by Barberon et al. (2016) and (Naseer *et al*., 2012), where a complete lack of suberin in the CASP1::CDEF1 mutant was documented. In spite of this confusion, we assume that the CASP1::CDEF1 in fact lack suberin in roots and that our observations support that suberin restricts leakage of K from the stele.

We observed that the mature parts of both nodal and seminal roots, where most suberin is deposited, were able to secure K within the stele during translocation (Fig. 5a & 5b; Fig. 6a & 6b). On the contrary, closer to the tip, where both root types deposited less or remained unsuberized (Fig. 1), both seminal and nodal roots lost K from the stele during translocation (Fig. 5c and D; Fig. 6c & 6d). In line with the results from Arabidopsis suberin mutants (Barberon *et al*., 2016), these observations support that suberin facilitate net uptake of K by preventing loss of K from the stele by sealing the vasculature during translocation towards the shoot.

### Passage cells may prevent K loss

Some parts of the suberin barrier remains incomplete also in the mature part of roots due to the presence of unsuberized passage cells (Fig. 1) (Geldner, 2013; Holbein *et al*., 2021). Data implies that passage cells have multiple roles in root functionality, including serving specialized roles in ion transport. Thus, passage cells are always associated with the xylem poles and high expression of phosphate transporters, including PHO1, has been observed in passage cells and associated cortex cells in mature parts of Arabidopsis roots (Andersen et al. 2018).

There are striking differences in nodal and seminal root morphology, yet their respective functional roles in nutrient uptake remains unresolved. Nodal roots have larger diameters than seminal roots, which translates into a larger xylem area, and it has been shown that they have a higher capacity for NO_3_^-^ and Ca uptake (Robards *et al*., 1973; Liu *et al*., 2020). We observed that nodal roots have similar capacity for K uptake as seminal roots when quantifying the resulting ^41^K enrichment in leaves (Fig. 2). This is in agreement with data obtained by Liu et al. (2020) showing an equal accumulation of the K-tracer Rb in shoots from nodal and seminal roots in a split root system (Liu et al., 2020). Accordingly, we observed no K leakage from the stele of mature sections of barley nodal roots, albeit suberin only deposits in a phloem-facing pattern here, thus leaving multiple passage cells unsuberized (Fig. 1; Fig. 6a & 6b). Several ion transporters appear to be expressed specifically in passage cells and their associated cortex cells, including the outward K channel SKOR in Arabidopsis (Gaymard et al., 1998). Thus, it is likely that, while suberin restricts loss of K at the suberized endodermal cells facing the phloem, the activity of K transporters in passage cells and associated cells are likely to rescue K that would otherwise be lost (Fig. 7B).

**Figure 7.**
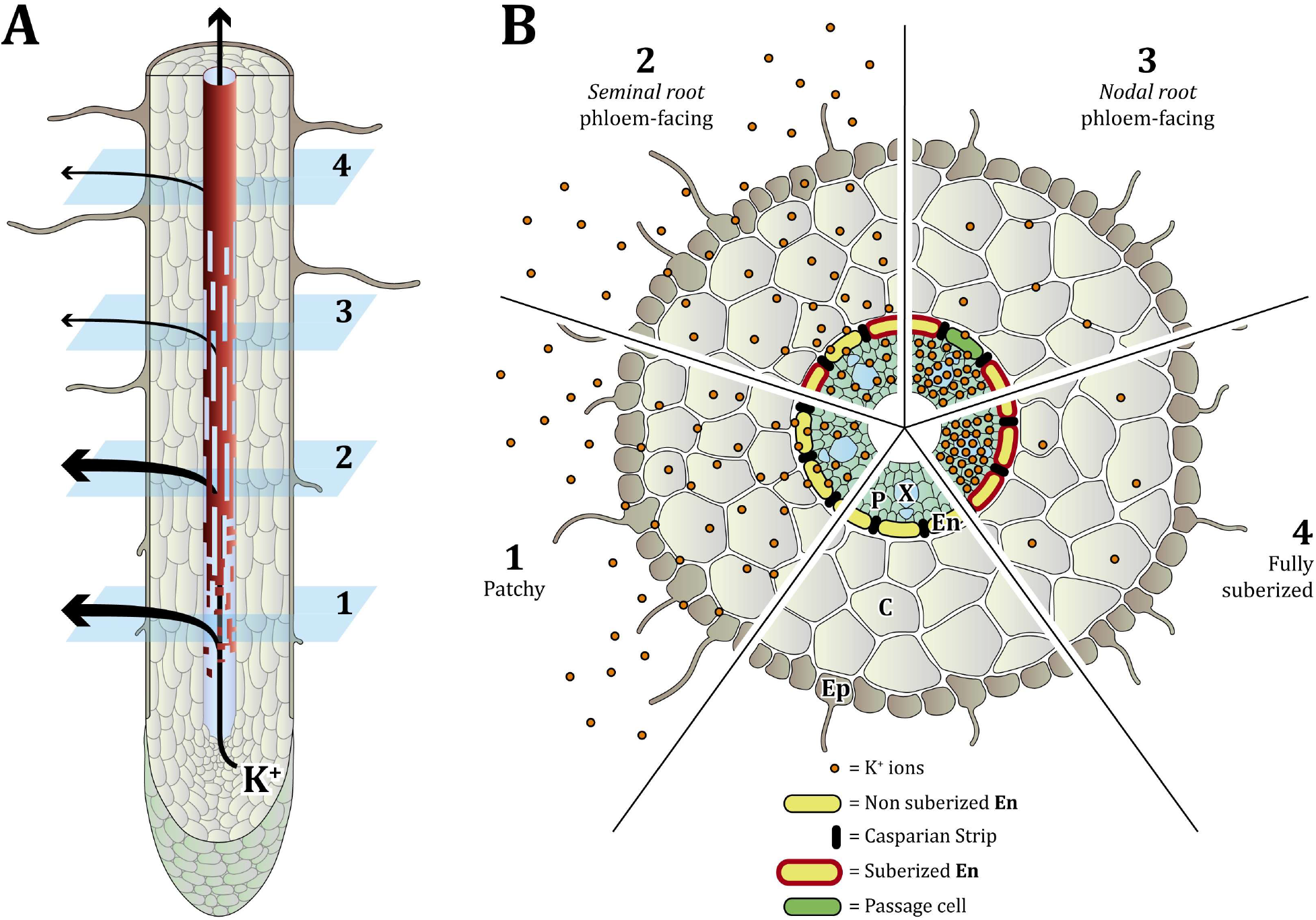
Radial outward transport of K from maturation zones differing in endodermal suberization. (1) suberin deposits in the secondary cell wall of a few scattered cells in the patchy zone. Following the patchy zone there is a phloem-facing zone (2) characterized by a deposition of suberin exclusively in cell walls of endodermal cells facing the phloem poles of the stele. In the mature part of the root, unsuberized cells in the phloem facing zone might adopt a passage cell identity (3). Towards the base of seminal roots, the suberin layer covers the endodermal cells fully (4). In the patchy (1) and immature phloem facing zone (2), K leaks out from the stele during translocation. In contrast, leakage of K across the endodermis is efficiently restricted in the mature phloem-facing zone (3) and the fully suberized zone (4). Ep, epidermis; C, cortex; En, endodermis; P, Phloem pole; X, Xylem.

Contrasting observations were made in the phloem-facing zones of seminal and nodal roots sampled at different distances to the root apex (Fig. 5 and 6). However, endodermal cells undergo a sequence of maturations steps during root development, which can readily be observed in barley as Casparian strips, suberin and tertiary thickenings. These steps reflect molecular changes as the endodermis matures sequentially starting at cell division of cortical-endodermal initials in the root apical meristem (Seo et al., 2021). Although the developmental fate of passage cells is predetermined by their association to the xylem pole, their molecular composition and functional properties likely changes as the root matures. Samples representing phloem-facing suberin zones were taken from different maturation states of nodal and seminal roots, as sections of the phloem-facing zone was sampled at 25% and 75% distance from the root tip of seminal and nodal roots, respectively. Accordingly, the ability of seminal and nodal roots to reduce K leakage from their respective phloem-facing zone are likely to be influenced by differences in endodermal cell maturity (Fig. 7).

Since ion uptake is energetically costly, a transporter-mediated rescue of K, instead of physically sealing the endodermis with suberin, must yield some other advantages to the plant. Such advantages could be related to the ability of passage cells to allow radial uptake of various essential nutrients which would otherwise be restricted by suberin, as well as facilitating plant-rhizosphere communication via transfer of hormones and signaling compounds. Thus, passage cells should possibly be regarded as a hot-spots for controlled exchange of chemical compounds, including mineral ions. (Barberon et al. 2016; Chen et al. 2019; Robbins et al. 2014).

In conclusion, this study provides experimental data supporting the role of suberin in restricting loss of K from the stele upon root to shoot translocation, and it provides ionomic data that suggest a crucial role for passage cells in reducing K leakage across the endodermis. However, owing to the single cell nature of passage cells, and lack of suitable analytical techniques with cellular resolution, it is not possible to provide further functional evidence for the role of passage cells in preventing ion leakage at present. Confirming these findings, will only be possible with further advances in single cell ionomics and transcriptomics technologies (Zhang et al., 2019; Holbein et al., 2021). In this future work, nodal roots will provide an excellent subject of study, because of their abundance of easy detectable passage cells and due to their documented role in limiting K leakage in the present work.

## ACKNOWLEDGEMENTS

We acknowledge the excellent technical assistance of Lena Byrgessen and Thomas Hesselhøj Hansen, and we are grateful for the financial support we received from the Danish Research council; Nature and universe (FNU).

## AUTHOR CONTRIBUTIONS

MVW, DAP and SHU designed the research. MWV and DAP performed the research. MWV, DAP and SHU analysed the data. MWV drafted the manuscript. DAP and SHU revised the manuscript. FM illustrated figure 7.

## DATA AVAILABILITY

Data that support the findings in this study are available from the corresponding author upon request.

## REFERENCES

Barberon M, Geldner N. 2014. Radial Transport of Nutrients: The Plant Root as a Polarized Epithelium. Plant Physiology 166: 528–537.

Barberon M, Vermeer JEMEM, De Bellis D, Wang P, Naseer S, Andersen TGG, Humbel BMM, Nawrath C, Takano J, Salt DEE, et al. 2016. Adaptation of Root Function by Nutrient-Induced Plasticity of Endodermal Differentiation. Cell 164: 447–459.

Baxter I, Hosmani PS, Rus A, Lahner B, Borevitz JO, Muthukumar B, Mickelbart M V., Schreiber L, Franke RB, Salt DE. 2009. Root suberin forms an extracellular barrier that affects water relations and mineral nutrition in Arabidopsis (GP Copenhaver, Ed.). PLoS Genetics 5: e1000492.

De Bellis D, Kalmbach L, Marhavy P, Daraspe J, Geldner N, Barberon M. 2022. Extracellular vesiculo-tubular structures associated with suberin deposition in plant cell walls. Nature Communications 13.

Calvo-Polanco M, Ribeyre Z, Dauzat M, Reyt G, Hidalgo-Shrestha C, Diehl P, Frenger M, Simonneau T, Muller B, Salt DE, et al. 2021. Physiological roles of Casparian strips and suberin in the transport of water and solutes. New Phytologist 232: 2295–2307.

Che J, Yamaji N, Ma JF. 2018. Efficient and flexible uptake system for mineral elements in plants. New Phytologist 219: 513–517.

Chen A, Husted S, Salt DE, Schjoerring JK, Persson DP. 2019. The Intensity of Manganese Deficiency Strongly Affects Root Endodermal Suberization and Ion Homeostasis. Plant physiology 181: 729–742.

Cohen H, Fedyuk V, Wang C, Wu S, Aharoni A. 2020. SUBERMAN regulates developmental suberization of the Arabidopsis root endodermis. The Plant Journal: tpj.14711.

Doblas VG, Geldner N, Barberon M. 2017. The endodermis, a tightly controlled barrier for nutrients. Current Opinion in Plant Biology 39: 136–143.

Enstone DE, Peterson CA, Ma F. 2002. Root Endodermis and Exodermis: Structure, Function, and Responses to the Environment. Journal of Plant Growth Regulation 21: 335–351.

Farooq M, Wahid A, Kobayashi N, Fujita D, Basra SMA. 2009. Plant drought stress: Effects, mechanisms and management. Agronomy for Sustainable Development 29: 185–212.

Franke RB, Dombrink I, Schreiber L. 2012. Suberin goes genomics: use of a short living plant to investigate a long lasting polymer. Frontiers in plant science 3: 4.

Gaymard F, Pilot G, Lacombe B, Bouchez D, Bruneau D, Boucherez J, Michaux-Ferrière N, Thibaud JB, Sentenac H. 1998. Identification and disruption of a plant shaker-like outward channel involved in K+ release into the xylem sap. Cell 94: 647–655.

Geldner N. 2013. The Endodermis. Annual Review of Plant Biology 64: 531–558.

Grube Andersen T, Naseer S, Ursache R, Wybouw B, Smet W, De Rybel B, Vermeer JEM, Geldner N. 2018. Diffusible repression of cytokinin signalling produces endodermal symmetry and passage cells. Nature Publishing Group 555.

Hampton CR, Bowen HC, Broadley MR, Hammond JP, Mead A, Payne KA, Pritchard J, White PJ. 2004. Cesium toxicity in Arabidopsis. Plant physiology 136: 3824–3837.

Hetherington AJ, Dolan L. 2018. Stepwise and independent origins of roots among land plants. Nature 561: 235–238.

Holbein J, Franke RB, Marhavý P, Fujita S, Górecka M, Sobczak M, Geldner N, Schreiber L, Grundler FMW, Siddique S. 2019. Root endodermal barrier system contributes to defence against plant-parasitic cyst and root-knot nematodes. Plant Journal 100: 221–236.

Holbein J, Shen D, Andersen TG. 2021. The endodermal passage cell – just another brick in the wall? New Phytologist 230: 1321–1328.

Kawamoto T, Kawamoto K. 2014. Preparation of thin frozen sections from nonfixed and undecalcified hard tissues using Kawamot’s film method (2012). Methods in molecular biology (Clifton, N.J.) 1130: 149–164.

Kopittke PM, Lombi E, van der Ent A, Wang P, Laird JS, Moore KL, Persson DP, Husted S. 2020. Methods to visualize elements in plants. Plant Physiology 182: 1869–1882.

Kreszies T, Shellakkutti N, Osthoff A, Yu P, Baldauf JA, Zeisler-Diehl V V., Ranathunge K, Hochholdinger F, Schreiber L. 2018. Osmotic stress enhances suberization of apoplastic barriers in barley seminal roots: Analysis of chemical, transcriptomic and physiological responses. New Phytologist.

Krishnamurthy P, Ranathunge K, Franke R, Prakash HS, Schreiber L, Mathew MK. 2009. The role of root apoplastic transport barriers in salt tolerance of rice (Oryza sativa L.). Planta 230: 119–134.

Li B, Kamiya T, Kalmbach L, Yamagami M, Yamaguchi K, Shigenobu S, Sawa S, Danku JMC, Salt DE, Geldner N, et al. 2017. Role of LOTR1 in Nutrient Transport through Organization of Spatial Distribution of Root Endodermal Barriers. Current Biology 27: 758–765.

Lilay GH, Castro PH, Campilho A, Assunção AGL. 2019. The Arabidopsis bZIP19 and bZIP23 Activity Requires Zinc Deficiency – Insight on Regulation From Complementation Lines. Frontiers in Plant Science 0: 1955.

Liu Z, Giehl RFH, Hartmann A, Hajirezaei MR, Carpentier S, von Wirén N. 2020. Seminal and Nodal Roots of Barley Differ in Anatomy, Proteome and Nitrate Uptake Capacity. Plant and Cell Physiology 61: 1297–1308.

Middleton LJ, Handley R, Overstreet R. 1960. Relative Uptake and Translocation of Potassium and Cesium in Barley. Plant physiology 35: 913–8.

Naseer S, Lee Y, Lapierre C, Franke R, Nawrath C, Geldner N. 2012. Casparian strip diffusion barrier in Arabidopsis is made of a lignin polymer without suberin. Proceedings of the National Academy of Sciences of the United States of America 109: 10101–10106.

Palmgren M. 2018. Plant epithelia: What is the role of the mortar in the wall? 16: e3000073.

Persson DP, Chen A, Aarts MGM, Salt DE, Schjoerring JK, Husted S. 2016. Multi-Element Bioimaging of Arabidopsis thaliana Roots. Plant physiology 172: 835–847.

Robards AW, Jackson SM, Clarkson DT, Sanderson J. 1973. The structure of barley roots in relation to the transport of ions into the stele. Protoplasma 77: 291–311.

Robbins NE, Trontin C, Duan L, Dinneny JR. 2014. Beyond the barrier: Communication in the root through the endodermis. Plant Physiology 166: 551–559.

Seo DH, Jeong H, Choi Y Do, Jang G. 2021. Auxin controls the division of root endodermal cells. Plant Physiology 187: 1577–1586.

Shukla V, Han JP, Cléard F, Lefebvre-Legendre L, Gully K, Flis P, Berhin A, Andersen TG, Salt DE, Nawrath C, et al. 2021. Suberin plasticity to developmental and exogenous cues is regulated by a set of MYB transcription factors. Proceedings of the National Academy of Sciences of the United States of America 118.

Ursache R, De Jesus Vieira Teixeira C, Dénervaud Tendon V, Gully K, De Bellis D, Schmid-Siegert E, Grube Andersen T, Shekhar V, Calderon S, Pradervand S, et al. 2021. GDSL-domain proteins have key roles in suberin polymerization and degradation. Nature Plants.

Zhang TQ, Xu ZG, Shang GD, Wang JW. 2019. A Single-Cell RNA Sequencing Profiles the Developmental Landscape of Arabidopsis Root. Molecular Plant 12: 648–660.

